# Artefactual formation of pyruvate from in-source conversion of lactate

**DOI:** 10.1101/256271

**Authors:** Sophie Trefely, Clementina Mesaros, Pening Xu, Mary T. Doan, Helen Jiang, Oya Altinok, Zulfiya Orynbayeva, Nathaniel W. Snyder

## Abstract

**Rationale:** Lactate and pyruvate are high abundance products of glucose metabolism. Analysis of both molecules as part metabolomics studies in cellular metabolism and physiology have been aided by advances in liquid chromatography-mass spectrometry (LC-MS).

**Methods:** We used ion pairing-chromatography and negative ion mode ESI on an QExactive HF to perform stable isotope assisted metabolomics profiling of lactate and pyruvate metabolism.

**Results:** Using an LC-MS method for polar metabolite analysis we discovered an artefactual formation of pyruvate from in-source fragmentation of lactate. Surprisingly, this in-source fragmentation has not been previously described, thus we report this identification to warn other investigators. This artefact was detected by baseline chromatographic resolution of lactate and pyruvate by LC with confirmation of this artefact by stable isotope labeling of lactate and pyruvate.

**Conclusions:** These findings have immediate implications for metabolomics studies by LC-MS and direct infusion MS, especially in negative ion mode, whereby users should resolve lactate from pyruvate or robustly quantify the potential formation of pyruvate from higher abundance lactate in their assays.

## Introduction

Lactate and pyruvate are interdependent major three carbon products of glucose metabolism. Due to the different functions and routes to generate both lactate and pyruvate, measurement of their abundance, metabolic source, and fate remains analytically important(1). Concentrations of lactate in different matrices and under different conditions can range widely, as can levels of pyruvate. Blood lactate can transit from resting 1 mM to 20 mM within seconds of aerobic exercise. Lactate is at considerably higher concentrations than pyruvate in most bioanalytical samples, including blood at CSF (1.3 mM and 1.5 mM) with pyruvate at (0.124 mM and 0.110 mM) in children(2). Lactate to pyruvate ratios can be indicative of metabolic dysfunction, with high lacate:pyruvate indicating respiratory chain, Krebs cycle, and pyruvate carboxylase defects and with low lacate:pyruvate ratios indicating pyruvate dehydrogenases defects. Thus, dynamic range in clinical and experimental measurement of lactate is critical, with specificity, sensitivity and dynamic range being of varying importance in measurement of pyruvate. Common assays for lactate utilize enzymatic conversion of lactate via lactate oxidase or lactate dehydrogenase, and detect L-lactate. An alternate assay may be necessary for D-lactate, or in situations where interferences with lactate measurement (for example by glycolate) may occur(3). Studies employing mass spectrometry (MS), in particular high resolution mass spectrometry (HRMS), coupled to liquid chromatography (LC) or gas chromatography (GC) have increasingly gained prominence for their application to semi-quantitative, quantitative, and stable isotope tracing experiments. In particular, the wider adoption of standalone HRMS (commonly referred to as flow injection analysis (FIA) or direction infusion), and LC-HRMS, allows unique opportunities for faster and more sensitive analytical workflows. MS-based assays have been deployed to query remaining questions about the function, localization and context dependent metabolism of pyruvate and lactate.

Pyruvate can be shuttled across the mitochondrial membrane to form acetyl-CoA via the pyruvate dehydrogenase complex (PDC) and enter the Krebs cycle for oxidation. Defects in the pyruvate carrier result in metabolic effects on high sugar diets and clinical effects in affected families with varying intensity(4). Identification of the functional importance of PDC in the mammalian nucleus has also shown that pyruvate may directly contribute to nuclear acetyl-CoA pools and thus potentially histone acetylation(5). Alternatively, pyruvate can be metabolized to lactate anaerobically by cytosolic lactate dehydrogenase (EC 1.1.1.27) with concomitant regeneration of NAD+ from NADH. The equilibrium constant of lactate dehydrogenase couples cytosolic pyruvate and lactate, thus the pools of lactate and pyruvate are interdependent. In anaerobic settings, this resulting lactate is exported via monocarboxylate transporters and can reach high concentrations in extra-cellular fluids. Similarly, in ischemia of tissues, lactate can reach high intracellular concentrations and high concentrations within tissues. The formation of pyruvate from lactate occurs in normal physiology as part of the Cori cycle, with recent evidence that this process also occurs pathologically during cancer metabolism. Interest in lactate has been renewed by an increased understanding that lactate may serve as a critical carbon source in certain tumor and cell types, presumably by conversion to pyruvate and then incorporation into acetyl-CoA(6). The localization and mechanism of how lactate metabolism is rewired in different cells and tissues remains an active area of research(7). Thus, physiological measurements of lactate span importance in cellular metabolism, physiology, tumor metabolism, and blood chemistry.

During an LC-HRMS experiment for lactate and pyruvate quantification, an additional high intensity peak was noted in the chromatogram for pyruvate. Further investigation, including using stable isotope labeled analogs of lactate and glucose, revealed this artefactual peak to come from a previously unreported in-source formation of pyruvate from lactate.

## Materials and Methods

### Materials

Optima LC-MS grade water, methanol, and acetonitrile (ACN) were purchased from Thermo Fisher Scientific (Waltham, MA). [^13^C_6_]-Glucose, diisopropylethylamine (DIPEA), and 1,1,1,3,3,3-hexafluoro 2-propanol (HFIP) were purchased from Sigma-Aldrich (St Louis, MO). [^13^C_3_]-lactate was from Cambridge isotope laboratories (Tweksbury, MA).

### Cell line and growth conditions

Human colon cancer HCT116 cells (ATCC, Manassas, VA) were cultured for passage in RPMI1640 medium supplemented with 10% FBS at 37°C in 5% CO_2_ atmosphere.

### Isolation and isotopic labeling of mitochondria from cultured cells

Mitochondria were isolated by differential centrifugation. Approximately 1 *g* of cell pellet rinsed with ice-cold PBS (without Ca^+2^/Mg^+2^) was re-suspended in hypotonic solution (100 mM sucrose, 10 mM MOPS, pH 7.2, 1 mM EGTA) for 10 minutes (min) on ice to allow cells to swell. The suspension was homogenized in a Teflon-glass homogenizer and immediately isotonicity restored by addition of 100 μL hypertonic solution (1.25 M sucrose, 10 mM MOPS, pH 7.2) per each 1 mL of suspension. The mix was diluted with three volumes of isolation buffer (75 mM mannitol, 225 mM sucrose, 10 mM MOPS, pH 7.2, 1 mM EGTA, 0.1% fatty acid free BSA). Cellular debris were sediment at 980 × *g*, 4°C for 5 min. Supernatant containing mitochondria was centrifuged at 10,300 × *g*, 4°C, for 20 min. The resulting mitochondria pellet was rinsed by centrifugation at the same parameters in 10 mL of MiR06 respiration buffer (110 mM sucrose, 60 mM K-lactobionate, 20 mM HEPES, pH 7.2, 1 mM KH_2_PO_4_, 3 mM MgCl_2_×6H_2_O, 0.5 mM EGTA, 20 mM taurine, 0.1% fatty acid free BSA). The final mitochondria pellet was re-suspended in 100 μL MiR06 by vortexing and stored on ice. The mitochondria protein was determined by Bradford Protein Assay. The 10 mM lactate (unlabeled or tracer) was added to 0.1 mg mitochondria protein/sample in MiR06 respiration buffer containing 2 mM malate and 2 mM ADP in total 200 μL volume. The mix was incubated at 37°C for 40 min. The reaction was stopped by adding 800 μL of -80°C methanol.

### Analysis of pyruvate and lactate by LC-HRMS

Isolated mitochondria from HCT116 cells in 80:20 (v/v) methanol:water at -80 °C were extracted with 15 0.5 s pulses with a handheld sonicator (Fisher) and centrifuged at 16,000 × *g* for 10 min at 4 °C to remove insoluble debris. The supernatant was evaporated to dryness under nitrogen gas, resuspended in 100 μL 5% (v/v) 5-sulfosalicylic acid in water, centrifuged again at 16,000 × *g* for 10 min at 4 °C, and 5 μL was injected for analysis by LC-HRMS with modifications as indicated from a previously described method(8). Briefly, lactate and pyruvate were quantified by single ion monitoring (SIM) on an Ultimate 3000 UHPLC coupled to a recently (within 48 hrs) calibrated Q Exactive HF (Thermo). The MS used a heated-ESI-II probe with a steel needle insert for high-flow applications used consistently at the B position in an IonMax Source Housing with all source gasses being high-purity nitrogen from Airgas.

### Data and Statistical analysis

Data analysis was conducted in XCalibur 3.0 Quan Browser and Tracefinder 4.1 (Thermo) and statistical analysis was conducted in Excel 2016 (Microsoft) and Graph Pad Prism 7 (GraphPad Software). Area under the curves (AUCs) were calculated for peak integration with a 5 part per million (ppm) mass window. Isotopologue enrichment was calculated as previously published(9).

## Results and Discussion

### Identification of a lactate-derived in-source fragmentation corresponding to pyruvate

Lactate and pyruvate was analyzed from mitochondrial extracts by LC-HRMS. Both lactate and pyruvate were retained on reversed phase chromatography using ion-pairing (**Fig. 1**), with a retention time (R*t*) of 4.8-5.1 min for lactate, and 5.3-5.6 min for pyruvate. Slight differences in retention time were observed between targeted single ion monitoring (SIM) and full scan (FS) data due to the duty cycle of the instrument acquisition method. Baseline resolution of lactate and pyruvate peaks was maintained in all experiments. Both lactate and pyruvate were detected as [M-H]^-^ ions, with *m/z* of the corresponding base peak (*m/z*, dppm observed-theoretical) for pyruvate (87.0087, 0) and lactate (89.0244, 0). An interfering peak in the pyruvate channel was observed at approximately 50% relative intensity of the pyruvate peak in both SIM and FS (Fig. 1A and B). The R*t* of this interfering peak corresponded to lactate, with peak areas for the ion corresponding to lactate 550-times higher than for the interfering peak in the pyruvate channel. This raised the possibility that artefactual formation of pyruvate from lactate was being observed from an in-source reaction. The reverse reaction (formation of lactate from pyruvate) was not observed, as there was no interfering peak detected in the lactate channel.

**Figure 1.**
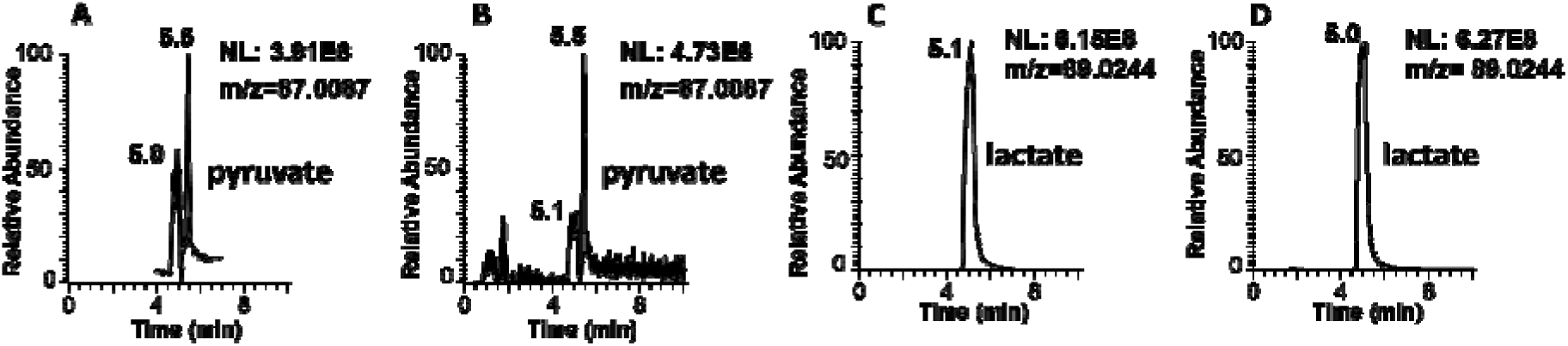
Liquid chromatography-high resolution mass spectrometry chromatograms for lactate and pyruvate using different scan modes. A. Pyruvate single ion monitoring (SIM), B. pyruvate full scan (FS), C lactate SI M, and D. lactate FS.

To further probe this phenomenon, we examined the chromatograms from an experiment using ^13^C_3_-lactate as a stable isotope tracer. ^13^C_3_-isotopologues of lactate and pyruvate were both detected as the [M-H]^-^ ions, with *m/z* of the corresponding base peak (*m/z*, dppm observed-theoretical) for pyruvate (90.0188, 0) and lactate (92.0345, 0) This resulted in an enrichment in the signal corresponding to ^13^C_3_-lactate, with a concomitant increase in the signal for [^13^C_3_]-pyruvate. Again, an interfering peak was observed in the ^13^C_3_-pyruvate channel at the retention time corresponding to [^13^C_3_]-lactate with a similar relative signal intensity to the lactate peak (**Fig. 2**). All three labeled carbon atoms were maintained in the artefact peak, indicating that the in-source reaction was not from an insource adduct with a carbon containing molecule which would be unlabeled. Notably, none of the mass windows for peak integration at 5 ppm overlapped for any pairs of these isotopologues, further eliminating non-specific overlap of mass windows as a possibility of ion signal (**Table 1**).

**Figure 2.**
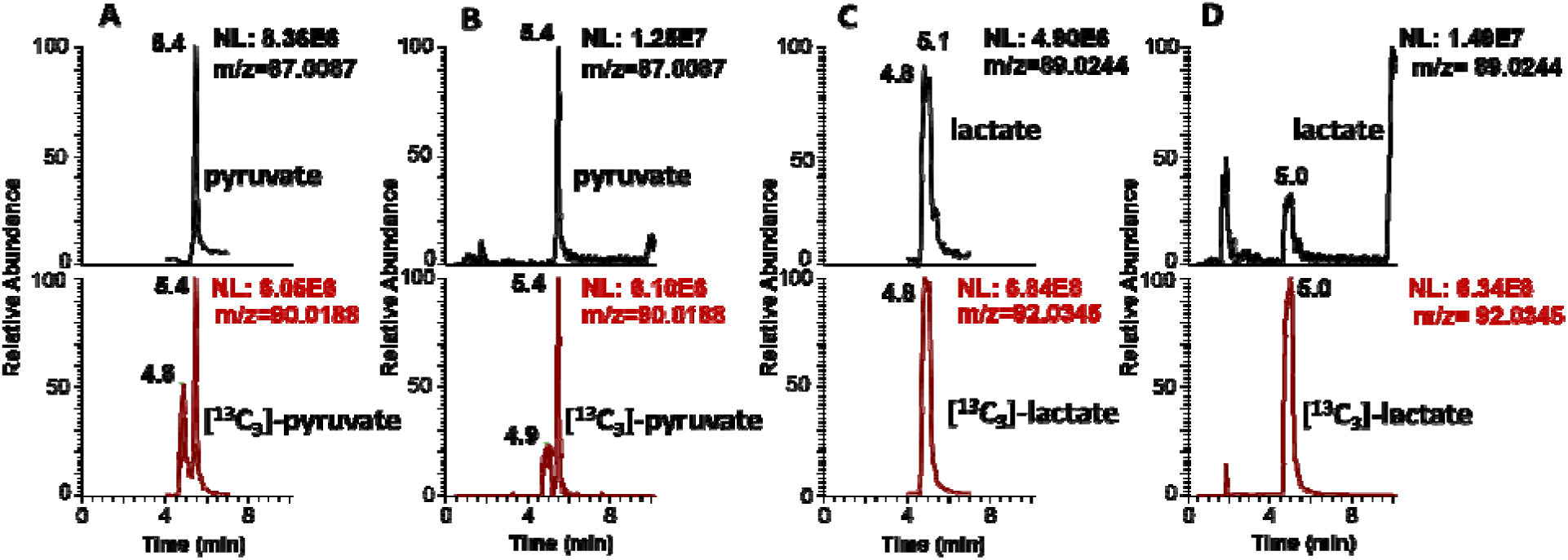
Liquid chromatography-high resolution mass spectrometry chromatograms for unlabeled lactate and pyruvate (top), and [^13^C_3_]-lactate and pyruvate (bottom), from isolated mitochondria incubated with [^13^C_3_]-lactate, using different scan modes from. A. Pyruvate single ion monitoring (SIM), B. pyruvate full scan (FS), C. lactate SIM, and D. lactate FS.

**Table 1.** Theoretical m/z values for pyruvate and lactate [M-H]^-^ with min and max m/z values for a 5 ppm mass window.

| Lactate | $[C_3H_5O_3]^-$ | $^{13}C_1$ | $^{13}C_2$ | $^{13}C_3$ |
| --- | --- | --- | --- | --- |
| m/z | 89.0244 | 90.0278 | 91.0311 | 92.0345 |
| m/z min | 89.0240 | 90.0273 | 91.0306 | 92.0340 |
| m/z max | 98.0249 | 90.0283 | 91.0316 | 92.0350 |
| Pyruvate | $[C_3H_3O_3]^-$ | $^{13}C_1$ | $^{13}C_2$ | $^{13}C_3$ |
| m/z | 87.0087 | 88.0121 | 89.0155 | 90.0188 |
| m/z min | 87.0083 | 88.0117 | 89.0151 | 90.0184 |
| m/z max | 87.0092 | 88.0125 | 89.0160 | 90.0193 |

In order to address the generalizability of this phenomenon and modifiable source parameters, we conducted experiments varying the spray voltage under the original solvent conditions. Increasing spray voltage from 2 to 4 kV increased lactate signal intensity, with a further increase to 4.5 kV reducing signal intensity (**Fig 3A**). The area under the curve for the in-source generated pyruvate generally increased with lactate, and plotting the ratios of pyruvate/lactate with varying spray voltage demonstrated an increase in pyruvate (**Fig 3A and B**).

**Figure 3.**
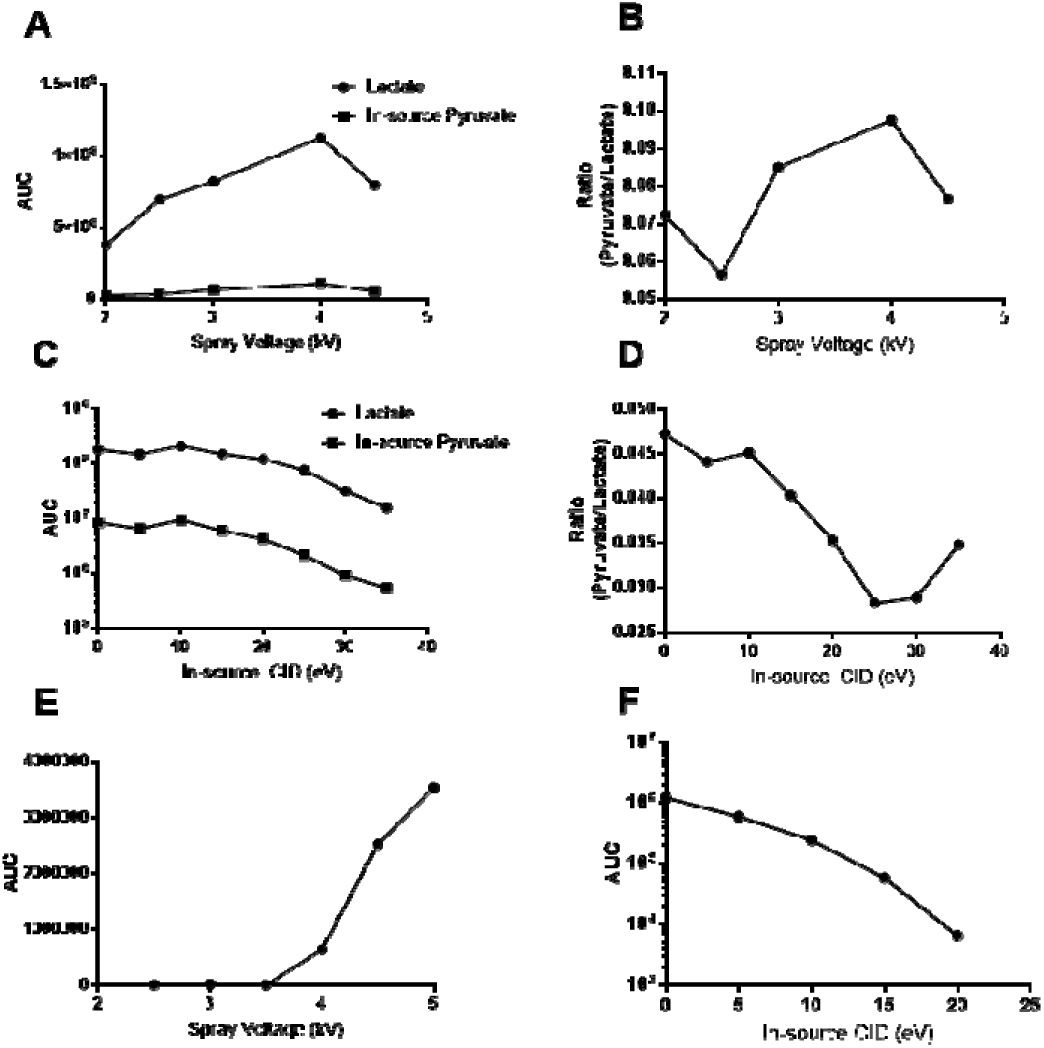
Source parameter influence on in-source generation of [^13^C_3_]-pyruvate from [^13^C_3_]-lactate. Varying spray voltage in negative ion mode for A. Area under the curve (AUC) of [^13^C_3_]-lactate and resulting [^13^C_3_]-pyruvate, and B. ratio of [^13^C_3_]-pyruvate/ [^13^C_3_]-lactate AUC. Varying in-source CID for C. Area under the curve (AUC) of [^13^C_3_]-lactate and resulting [^13^C_3_]-pyruvate, and D. ratio of [^13^C_3_]-pyruvate/ [^13^C_3_]-lactate AUC. AUC for [^13^C_3_]-lactate in positive ion mode with varying E. Spray voltage and F. In-source CID.

Next, we conducted direct infusion experiments using ^13^C_3_-lactate to reduce signal-to-noise, avoid contaminating our mass spectrometer, and test positive ion mode. ^13^C_3_-lactate was infused at 2 μL/min at 100 ng/μL in 90/10 water/methanol (v/v). Spray voltage was varied in positive ion mode, and in-source collision induced association was varied between 0 and 30 kV in both negative and positive ion mode. Spectra was collected within an 80-95 *m/z* SIM window for at least 50 scans, and ion intensity was extracted and plotted. In negative ion mode varying in-source CID did not decrease the lactate or pyruvate signal until above 10 eV, at which point both lactate and pyruvate intensity decreased, with pyruvate decreasing more rapidly until 30 eV. Unsurprisingly, using positive ion mode resulted in a less intense signal for lactate (e6 in positive vs e8 in negative for similar infusion). We did not detect an ion corresponding to the in-source generation of ^13^C_3_-pyruvate from the infused ^13^C_3_-lactate, even at higher spray voltage and lower in-source CID where ^13^C_3_-lactate was detected as an intense signal (Fig 3E and F). This data suggests that the in-source formation of pyruvate is specific to (or at least considerably more efficient in) negative ion mode. In comparison to the LC-HRMS experiments, the direct infusion solvent did not contain ion-pairing reagents (DIPEA and HFIP), but still used a water/methanol solvent.

Degradation and transformation of metabolites within the ion source can result in quantitative and qualitative problems, specifically, in complicating interpretation of spectra, reduced sensitivity, and difficulty studying degradation of the analyte in samples (10). For the more structurally complicated (compared to lactate and pyruvate) neonicotinoids, Chai, *et al*., described distinct modes of decomposition, with in-source fragmentation occurring in ESI. In-source reactions have been observed for other small polar metabolites, most notably glutamine and glutamic acid (11). Mono-, di-, and tri-phosphate and other common biochemical moieties have also been reported to generate problematic in-source fragmentation occurring in ESI. In-source reactions have been observed for other small polar metabolites, most notably glutamine and glutamic acid (11). Mono-, di-, and tri-phosphate and other common biochemical moieties have also been reported to generate problematic in-source fragmentation (12). Both Purwaha, *et al*., and Xu, *et al*., were able to observe and quantify this phenomenon due to resolution by LC, similar to our findings here. Like our findings, the in-source reaction of stable isotope labeled analytes was similar in fraction of peak intensity, thus allowing stable isotope labeled standards to correct for quantitative bias.

The potential mechanistic basis and implications of the in-source generation of pyruvate is worth note. Redox chemistry within an ESI source, as noted by John Fenn, “was too obvious to mention.” (13) However, the exact mechanisms, implications, and analytical impact of electrochemical processes within the ESI source are contentious even to experts (14). It is also non-trivial to exclude the potential for chemical electron transfer reactions in the liquid or gas phase (15). Previous literature on advantageous and adverse in-source electrochemical ionization has included oxidation of metal organic-complexes and aromatic systems and reduction of quinones. This work adds to the structural diversity of potential in-source reactants. Our system, like most commercial systems, uses a floated emitter with an upstream ground, so theoretically, both oxidation and reduction reactions are possible in the solvent system(13). This complication is a major caveat of our study, but may spur experiments with this phenomenon in more experimentally tractable sources. The totality of our available data suggests that the in-source generation of pyruvate from lactate is either a product of chemical electron transfer reaction or an electrochemical process in the ES source. This is supported by the observed specificity for negative ion mode for the oxidation of lactate to pyruvate but complicated by source designs available in our laboratory. We did not exhaustively test the effect of source type, matrix, solvents, source conditions, or instrument-specific parameters on this phenomenon since these will vary across metabolomics users. The immediate implications of this work is that users should investigate in their specific experimental context the formation of pyruvate from lactate as part of assay and method development.

## Conclusion

Resolution by ion-pairing chromatography of lactate and pyruvate, coupled with stable isotope labeling allowed identification of a previously unreported in-source formation of pyruvate from lactate. Due to the potential wide concentration difference in lactate and pyruvate in biological samples, analysts should ensure artefacts, even those resulting from low efficiency processes as observed here, do not interfere with accurate measurement of these analytes.

